# Random connectivity generates inhibitory microcircuits that decorrelate adaptation in visual cortex

**DOI:** 10.1101/2024.12.13.628375

**Authors:** Y. Kosiachkin, A. J. Hinojosa, S.E. Dominiak, B. Evans, L. Lagnado

**Affiliations:** Sussex Neuroscience, School of Life Sciences, University of Sussex, Brighton BN1 9QG, UK

## Abstract

Inhibitory neurons are fundamental to sensory processing in the cortex but the rules governing their connections with excitatory neurons are unclear. Are pyramidal cells with different functions generated through specific or random connections? We used two-photon imaging, optogenetics and modelling to investigate opposing forms of adaptation in layer 2/3 of mouse visual cortex. We find that a slow modulatory drive acts differentially depending on the relative strength of inputs that individual pyramidal cells receive from parvalbumin-positive and somatostatin-positive interneurons. The number of depressing and sensitizing pyramidal cells could be explained quantitatively by the simplest connectivity rule – all inhbitory synapses made randomly. The functional heterogeneity of the pyramidal cell population therefore begins with general statistics of the connectome - interneurons are much sparser and connect with probabilities far less than one. The resulting “patchwork” of inhibitory microcircuits causes modulatory inputs to strongly decorrelate pyramidal cells on the behavioural time-scale of seconds.

## Introduction

The varied ways in which sensory inputs are processed in the cortex depends on the diversity of excitatory and inhibitory neurons and the circuits in which they interconnect^1–8^. Inhibitory neurons come in a variety of morphological and functional types^9,10^ so there is the potential for a number of such circuits to be built. We now need to understand how far inhibitory connections vary between individual PCs within a layer of cortex and, most crucially, how such variations might alter their functional ^11–13^.

The large majority of interneurons can be placed in one of three major classes based on expression of the peptides parvalbumin (PV), somatostatin (SST) or vasoactive intestinal polypeptide (VIP) ^12^. One view is that these deliver a *uniform* “blanket” of inhibition to PCs ^5,14–16^, while another is that there are *specific* rules that connect PCs into different microcircuits, thereby adjusting their function^3,13,17,18^. Such molecularly-determined connectivity rules can be seen to operate when comparing pyramidal cells in different layers of cortex^19^ or projecting from layer 5 to different long-range targets ^20,21^ but it is not known how far they can explain inhibitory circuits *within* layers. A third possibility is that distinguishable microcircuits arise *randomly* based on general structural features of the cortex. This idea is encapsulated by "Peters’ Rule", which proposes that the number of synaptic connections between two neurons is proportional to the geometric overlap of their axonal and dendritic arbors^8,22–25^ rather than more specific genetic or functional targeting ^25,26^.

Peters’ Rule has been described as a “pragmatic approximation to neural circuitry” with the potential to “provide a statistical summary of the connectome”^24^ and it has been used to interpret structural and connectomic data from a wide range of neural circuits, including the retina^27–30^, somatosensory cortex^31,32^, visual cortex^33^, frontal cortex^26^, hippocampus^34^, the olfactory and visual ^35,36^ systems of *Drosophila* and the high vocal center of song birds^37^. In the retina and cortex, a random connectivity rule provides a good quantitative description of the wiring of some types of neuron ^8,26,30,38^ but not all ^8,24,28–30,39^. Ultimately, the aim of connectomics data is to understand how signals flow in circuits, which raises a general question: Can Peters’ rule also provide a statistical summary of function?

Using a combination of *in vivo* imaging, optogenetics and modelling we investigated variations in inhbitory microcircuits in layer 2/3 of V1 and how these determine adaptation - a key functional property of PCs. Adaptation is a found in all sensory systems and acts to alter neuronal sensitivity according to the recent history of the input^40,41^. In V1 adaptation is strikingly heterogenous, to the extent that opposing forms occur simultaneously across the PC population: some PCs respond with high initial gain and then depress while others respond with low initial gain and then sensitize^42^. Optogenetic manipulations indicate that these funtional differences reflect variations in the balance between inhibitory inputs tp PCS, with SST inputs driving sensitization and PVs driving depression^42^.

We find that the distribution of adaptive responses in V1 can be explained by the simplest statistics of the local connectome, the densities of PV and SST interneurons and their *average* connection probabilities to PCs, together with the simplest connectivity rule – all synapses made randomly. Two statistics of the local connectome cause PCs to operate under a “patchwork blanket” of inhibition: PV and SST interneurons are much sparser than PCs and they connect to them with probabilities far less than one. The resulting variations in the balance between PV and SST inputs cause profound differences in the way individual PCs adapt in response to slower modulatory signals arriving from other cortical areas^43,44^. This study provides a functional demonstration of the consequences of Peters’ rule for cortical processing ^24^ – decorrelation of the PC population.

## Results

The profound variations in how PCs in V1 adapt to a visual stimulus are shown in Fig. 1: while some start responding with high gain and then depress others start responding with low gain and then sensitize (Fig. 1D-E). To obtain a quantitative understanding of how differences in connectivity might generate these opposing forms of adaptation we built a populational rate-based model based on the dynamics of PC, PV, SST and VIP neurons measured *in vivo* in response to a high-contrast stimulus (Fig. 1C, E-H). We begin by describing the construction of the model and the testing of its predictive abilities. We then use it to quantify the differences in inhibitory microcircuits that cause opposing forms of adaptation and ask whether these can be accounted for by a random connectivity rule.

**Figure 1.**
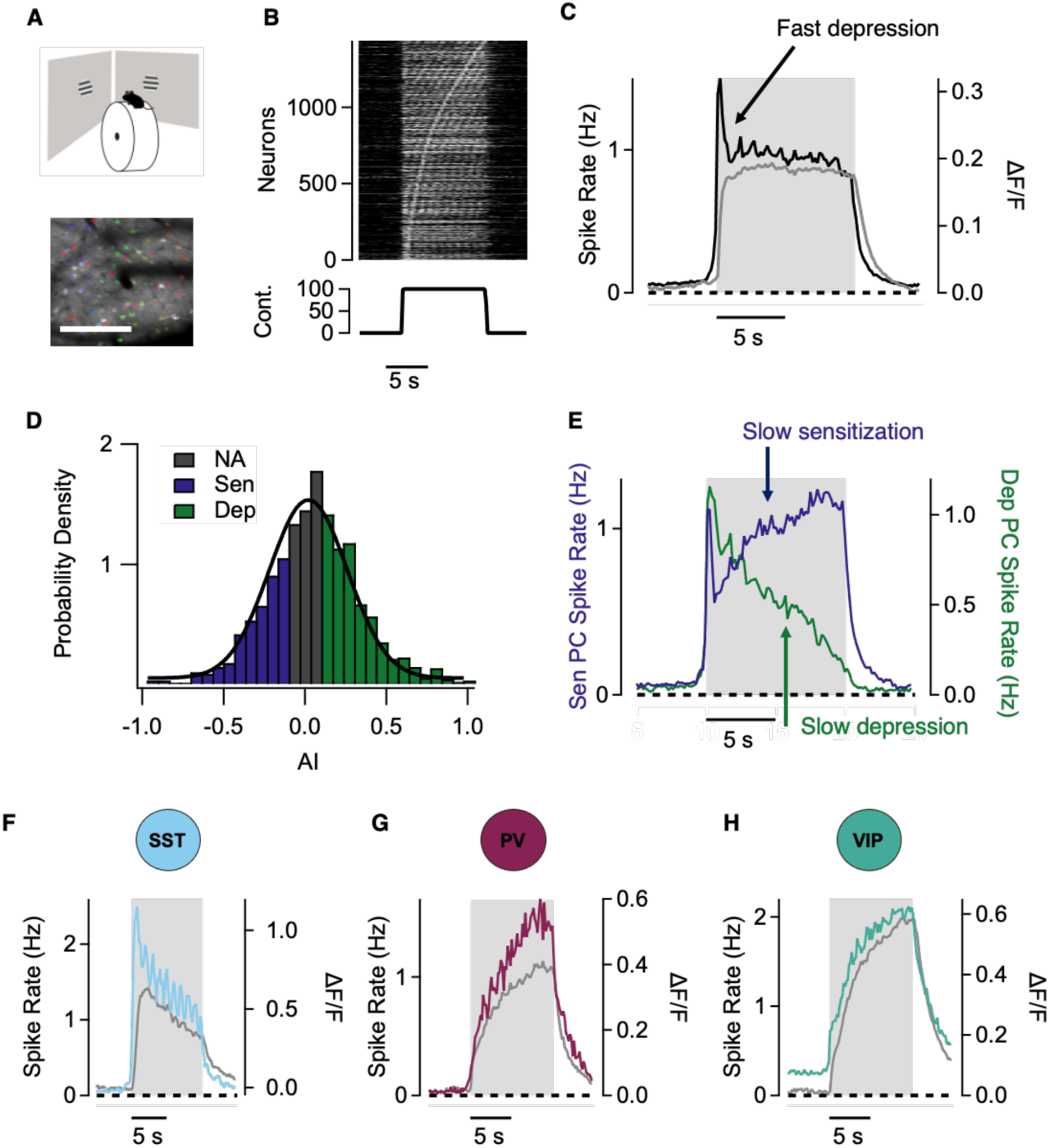
Slow adaptation in pyramidal cells and interneurons. **A**. Mice running on a treadmill were shown a visual stimulus (drifting grating, 20°) during 10 seconds (top). Example field of view (FOV) showing PCs in V1 expressing GCaMP6f (grey) and corresponding regions-of-interest (ROIs) for individual neurons (rainbow). Scale bar represents 200 µm. **B**. Raster plot showing deconvolved responses from PCs that were responsive to the stimulus (n = 1437 neurons, from 14 mice). **C**. Average response to the visual stimulus from calcium signals (grey, right axis) and deconvolved signal (black, left axis). Deconvolution reveals an initial peak of fast depression followed by a relatively flat slow response. **D**. Distribution of Adaptive index (AI) in the PC population in non- adaptors (grey), sensitizers (blue) and depressors (green). **E**. Splitting the distribution of PCs according to their AI reveals that the tertile with lower AI has slow sensitization (blue, right axis) while the tertile with higher AI has slow depression (green, left axis). **F, G and H.** Average population response to the visual stimulus in SSTs (n = 218, from 6 mice), PVs (n = 255, from 3 mice) and VIPs (n = 867, from 4 mice), respectively. In each case, the grey line is the GCaMP6f signal, and the coloured line is the inferred spike rate.

### A data-driven model of contrast adaptation

To construct a circuit model of adaptive processes in layer 2/3 we had to consider three general issues: *i)* the dynamics of activity in all the major types of neurons; *ii)* the dynamics of external signals that modulate activity during adaptation, and *iii)* the wiring of neurons within V1 as well as the inputs arriving from other regions of cortex.

***i) The dynamics of neurons in layer 2/3.*** Activity was measured by two-photon imaging of the calcium indicator GCaMP6f in mice running on a treadmill (Fig. 1A). Fast and slow phases of adaptation were observed in PCs responding to a drifting grating. The fast phase was not obvious in the raw GCaMP6f signal because of the relatively slow dynamics of the rise in bulk calcium concentration (Fig. 1B, C). We therefore used the Mlspike algorithm to infer spike rates from calcium signals ^45–47^ (see Methods). Fast adaptation in the inferred spike rate was dominated by depression and was complete within 1 s (Fig. 1C), as expected from electrophysiological recordings^48,49^. The slow phase of adaptation occurred over several seconds but varied widely across the PC population to the extent that while some neurons continued to depress others underwent strong sensitization (Fig. 1B, D, E).

A simple metric describing slow adaptation is the adaptive index (AI), calculated from the GCaMP6f signal as AI = (R1 − R2)/(R1 + R2) where R1 is the average response over the period 0.5- 2.5 s after stimulus onset and R2 is the average at 8-10 s. AI has a value of zero in the absence of slow adaptation, one when a neuron depresses completely and minus one when there is no initial response but the neuron then sensitizes (Fig. 1D). The functional heterogeneity across the PC population can be surveyed by ordering responses by AI (Fig. 1B) and then splitting the distribution into thirds (Fig. 1B and D). The average response in the first tertile clearly differentiates fast depression from a slow phase of sensitization (Fig. 1E, dark blue). The initial response was similar for PCs in the last tertile (green) after which slow adaptation occured in the opposite, depressing, direction (Fig. 1E). The general picture is that the process driving slow adaptation also decorrelates the PC population.

Adaptation was also an obvious feature of responses in the three main interneuron populations, but while SSTs were strongly depressing, PVs and VIPs were sensitizing ^42^ (Fig. 1F-H). The dynamics within each interneuron population were significantly more homogenous than the PC population (Supplementary Figure 1) and similar co-activation has been observed in the somatosensory cortex ^50^. The time-scales of slow adaptation were notably similar across all types of neuron, regardless of whether responses depressed or sensitized. For instance, the dynamics of depression were similar in SSTs and the subset of PCs where this form of adaptation dominated (Fig. 2A). Similarly, the dynamics of sensitization in VIPs and PVs were very similar to sensitization in PCs (Fig. 2B). A parsimonious explanantion for these observations is that a single signal dominates the time-course of depression and sensitization in different types of interneurons as well as the opposing forms of adaptation in PCs.

**Figure 2.**
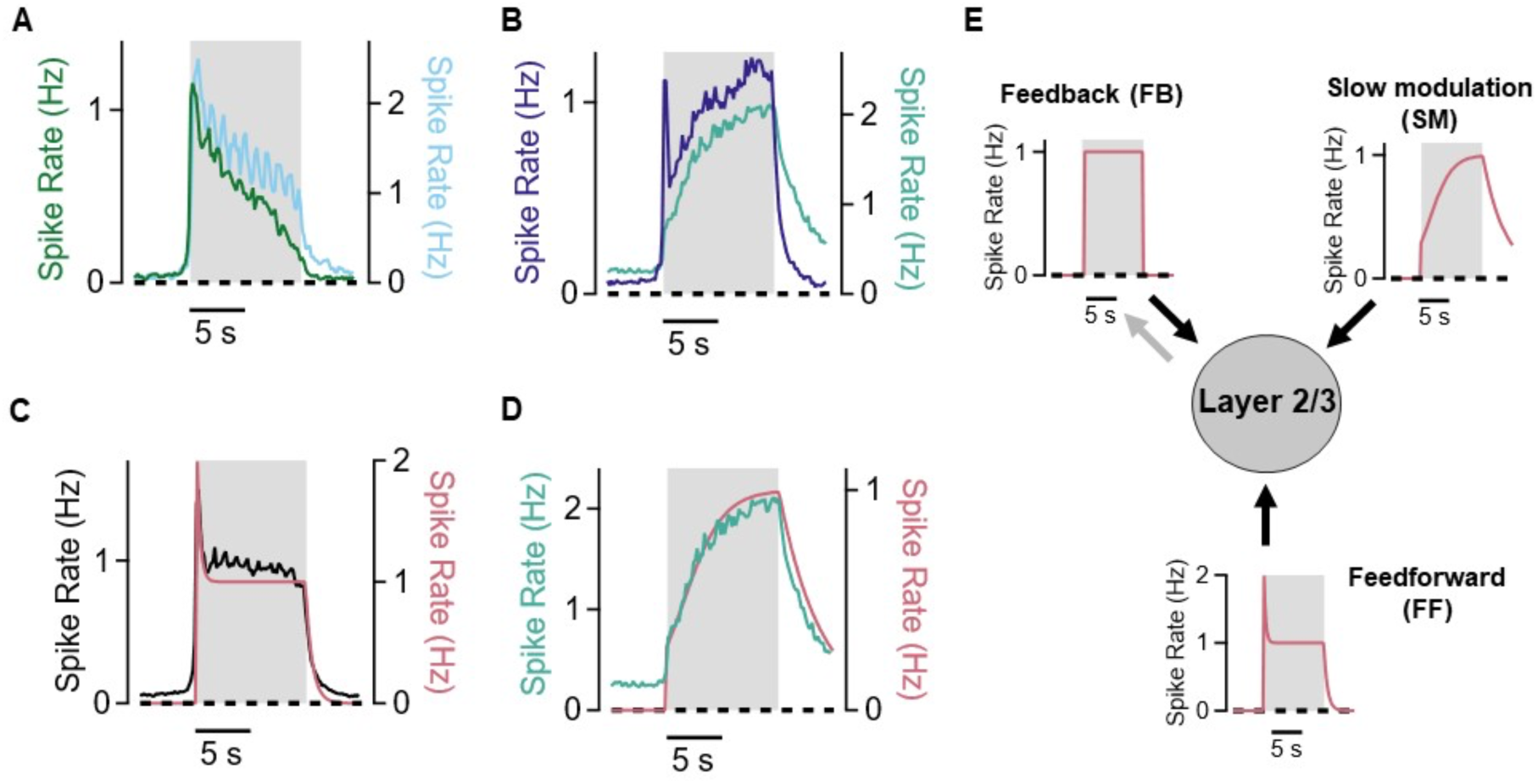
The dynamics of excitatory inputs to layer 2/3. **A**. Similar dynamics of slow depression in SSTs (blue, right axis) and depressing PCs (green, left axis). **B**. Similar dynamics of sensitization in VIPs (green, right axis) and in sensitizing PCs (blue, left axis). **C.** The feedforward input to the model (FF, red) accounted for fast depression and was based on the population average response of PCs (black). FF consists of a fast exponential decay with time-constant 0.21 s to a steady level 50% of the peak. **D**. The slow modulatory input to the model (SM, red) dominated the time-course of slow adaptation and was based on the average response of the VIP population (green) and consists of a step and a sigmoid function with a slow increase in activity (time-constant = 1.76 s). **E.** The three sources of external excitatory drive to layer 2/3. Feedback input (FB) was modelled simply as a step function with a delay of 380 ms. Scaling was chosen to roughly match the average activity of PCs.

***ii) The dynamics of signals entering layer 2/3.*** Excitatory signals entering layer 2/3 can be placed in three broad groups^51,52^: feedforward (FF) drive originating from the LGN, slower modulation from more distant cortical areas (SM) and feedback from higher visual areas (fb; Fig. 2) ^53–55^. FF inputs to layer 2/3 have long been known to be dominated by fast depressing adaptation occurring with time-constants of 300-600 ms^48,56^ and the fast phase of adaptation in PCs fell within this range (Fig. 1C). The average response across the PC population therefore served as the template for FF input (Fig. 2C).

V1 is also subject to slower “top-down” modulation from more distant cortical areas^54^. Inputs from the antero-medial region of frontal cortex, for instance, sensitize PC responses to a moving grating stimulus of the type used here and this occurs through adjustments in local inhibition^43^. Long-range inputs from the retrosplenial cortex also gradually ramp up over several seconds in response to a high-contrast stimulus during learning^57^. The temporal similarity suggests that such long-range inputs determine the common dynamics of slow adaptation across layer 2/3 (Fig. 2B and D). To test this idea we took advantage of the fact that a major target of long-range inputs are VIP interneurons which activate PCs through a VIP->SST->PC disinhibitory circuit, for instance during locomotion or changes in behavioural state ^44,58,59^. VIPs were inhibited optogenetically using ArchT while simultaneously measuring activity in SSTs. Control stimulus trials were interleaved with optogenetic trials and the responses averaged (Fig.3A, B). Slow depression in SSTs was almost completely abolished by suppressing activity in VIPs, indicating that these carried the SM signal generating adaptation in layer 2/3. We therefore used the average response of VIPs as the template for the SM input, which was described as the sum of a step and sigmoid function (Fig. 2D and E).

**Figure 3.**
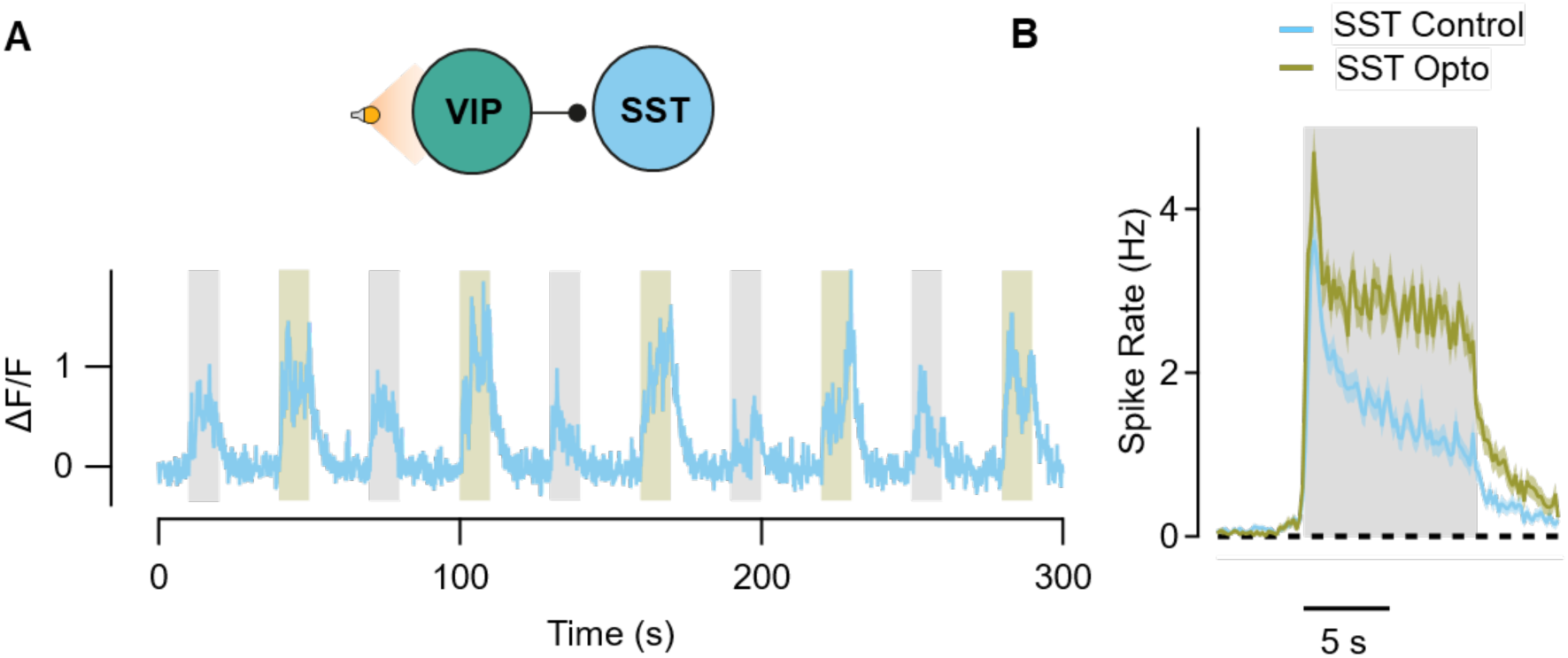
Slow adaptation in SST interneurons is driven by inputs from VIPs. **A**. Top: Scheme showing the experimental paradigm: To assess the role of VIP inhibition on SST dynamics we silenced VIPs while recording the activity of SST interneurons. Bottom: Example response of one SST neuron during stimulus presentation alone (grey bars) and stimulus paired with optogenetic silencing of VIPs (yellow). Silencing VIPs turns the slow dynamics of SST activity flat. **B.** Average response from SST interneurons during the presentation of the visual stimulus (blue) and when combined with optogenetic inactivation of VIP interneurons (green; n = 83, from 4 mice).

The third type of input to V1 is feedback from higher visual areas^43,60–63^ and SST interneurons are a major target for these, delaying their excitation^43,64,65^ (Supplementary Figure 2A). We have limited understanding of the dynamics of the FB signal over the time- scale of several seconds so for simplicity it was modelled as a step input beginning at a delay of 380 ms. Including this feedback allowed the model to account for the delay in the response of the SST population relative to other populations (Supplementary Figure 2B).

***iii) Connectivity within layer 2/3.*** Cortical connections have been probed using a number of approaches, including viral tracing ^66^, electrophysiology ^11,12,67–69^ and optogenetics ^68^. In constructing the model, we only considered connections that have been consistently observed in layer 2/3 and are strong enough to significantly influence the circuit, as summarized in Fig. 4A (see also Methods). Not all neuron types connect. For example, SST interneurns do not interact with each other while the very small number of connections from VIPs to PCs were neglected because these are not strong enough to significantly influence the circuit ^12,68,70^.

**Figure 4.**
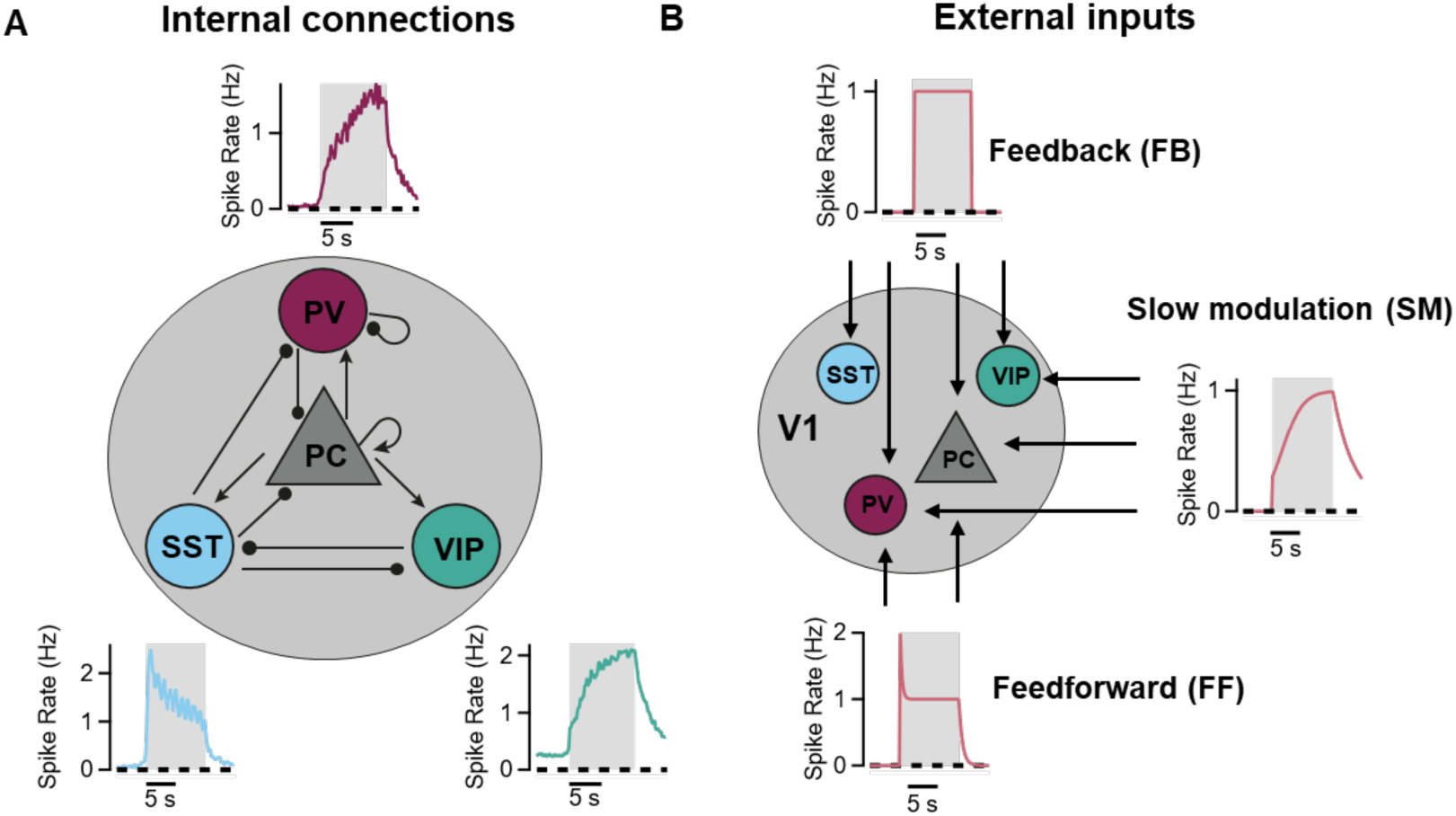
Connectivity within layer 2/3. **A.** The main excitatory and inhibitory connections between neural types within V1 include excitatory connections from PCs (arrows) and inhibitory connections from interneurons (round tip). **B.** Synaptic targets of external inputs. FF input targets PCs and PVs, FB input targets all neuron types and SM input targets PCs, PVs and VIPs.

The target neurons of the three types of input to layer 2/3 are shown in Fig. 4B. Feedforward excitation targets PVs and PCs^13^; feedback inputs targets all populations^71,72^ and the slower modulatory input enters through VIPs^59^ (Fig. 3) as well as directly activating PCs and PVs^73^. Definitive identification of the source of the SM signal is not required to use the model to estimate synaptic weights within layer 2/3 but likely candidates are the anteromedial and retrosplenial cortex^43,55,57^.

### Estimating synaptic weights in layer 2/3

The model used to estimate connection weights was based on first-order differential equations^74^. The *total* input from neuron population A to neuron population B is the average strength of the connection multiplied by the number of active A neurons, which we call here the total synaptic weight (see also Methods). In fitting the model, all synaptic weights were left free and varied to fit the average activity trace of each of the four cell populations (Fig. 5). The fit to the average response of the PCs, when there is very little slow adaptation, is shown in Fig. 5A. The fits closely followed both the fast and slow phases of the responses in PCs and all three interneuron types (Fig. 5A-D). The synaptic weights underlying these fits are shown in the heat map in Fig. 5E.

**Figure 5.**
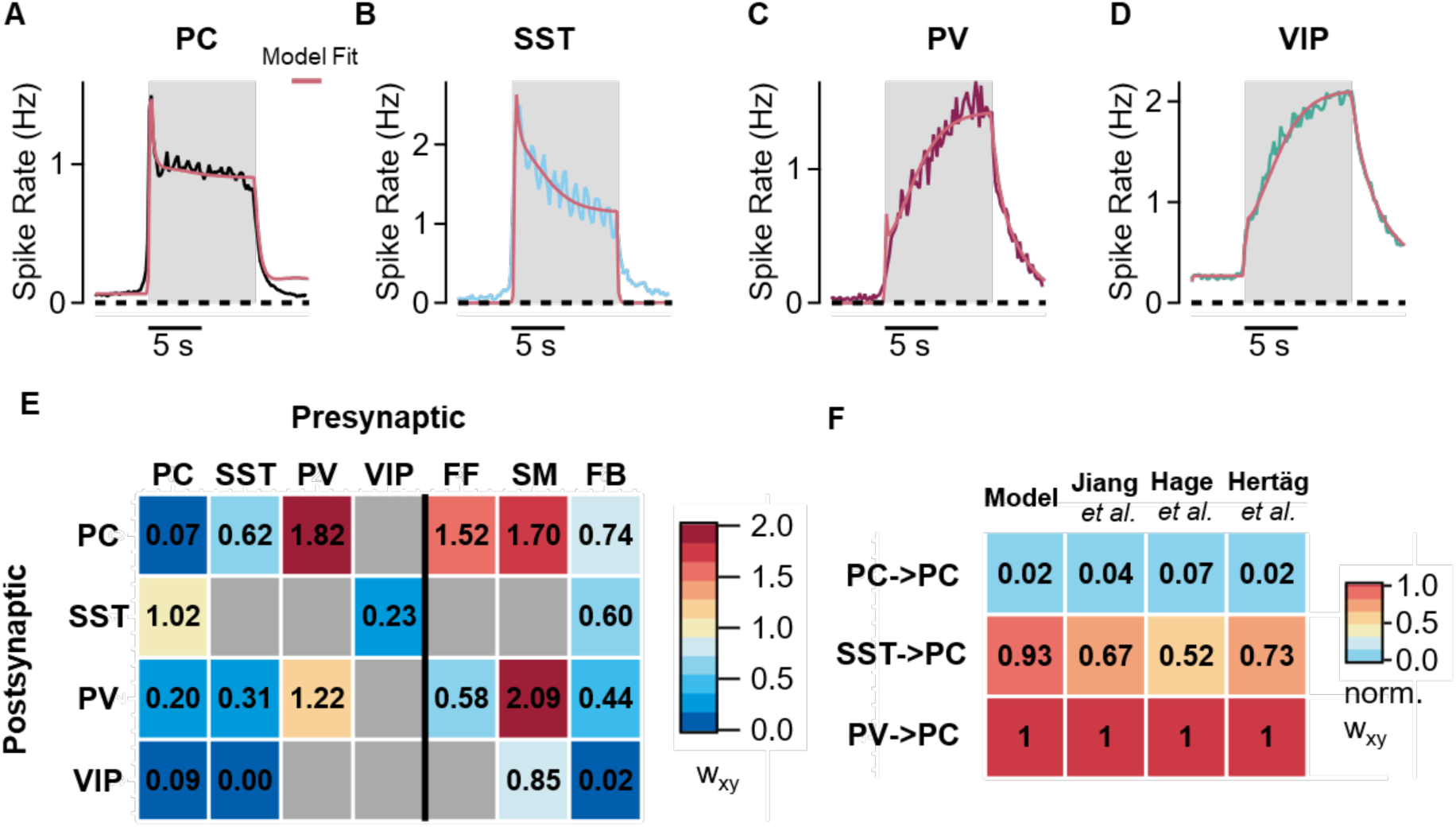
Synaptic weights predicting the dynamics of contrast adaptation in the four major neuronal populations in V1. **A-D**, Average firing rates of PCs (A, black), SSTs (B, blue), PVs (C, dark red) and VIPs (D, green) with their corresponding fits calculated by the model (light red). Note that these traces show the average activity of *responsive* neurons in each population. **E**, Heatmap of the synaptic weights (wxy) calculated by fitting the model to the activity of responsive neurons. The main excitatory input comes from external inputs, while inhibition is shared between SST and PV. Note that inhibition in the PC population is dominated by PV inputs. Grey boxes correspond to connections not considered in the model in E. **F**, Comparison of the relative synaptic weights received by PCs in the model compared to two studies using electrophysiology ^68,69^ and another modelling approach ^75^. Note that to allow comparison with these studies the synaptic weights in E were corrected according to the proportion of neurons in each population that responded to the stimulus.

The utlity of the model was first tested against established estimates of synaptic weights made using elelctrophysiology ^68,69^ as well as another model ^75^. To allow this comparison, we needed to account for the fact that our model was based on activity traces averaged only over the *responsive* neurons while these other studies measure the average connection strength between *complete* populations. The proportion of responsive neurons was, however, measured for each neuronal population, allowing a simple correction (see Methods). Fig. 5F shows the synaptic weights *averaged* across each population directly connecting PCs (all normalized to the weight of the strongest connection). There is a clear consensus that PCs receive their strongest inhibitory inputs from PV interneurons with inhibition from SSTs being weaker and excitation from other PCs negligible.

The model also predicted that feedback to VIPs is far weaker than feedback to SSTs and PVs (Fig. 5E), as has been observed experimentally for projections from the latero-medial area (LM) ^71^. There is less consensus for other synaptic weights such as, for instance, inputs to PVs, so we also took an experimental approach to testing the model more broadly. The activity of PV and SST interneurons was increased or decreased using optogenetics while monitoring activity in PCs. Fig. 6A (top) shows how the response of PCs was reduced when doubling the activity of SST interneurons using ChrimsonR ^42^. This decrease in gain was accurately predicted by the model simply by doubling the amplitude of responses in the SST population (red) while keeping all the other synaptic weights at the values obtained by fits to the control conditions (see Methods). Reducing the activity of SST interneurons by a factor of about ∼2 using ArchT had the opposite effect, amplifying the average PC response, which was also accurately accounted for (Fig. 6A, bottom). The model was similarly successful in predicting changes in response ampitude when optogenetically activating or inhibiting PVs (Fig. 6B).

**Figure 6.**
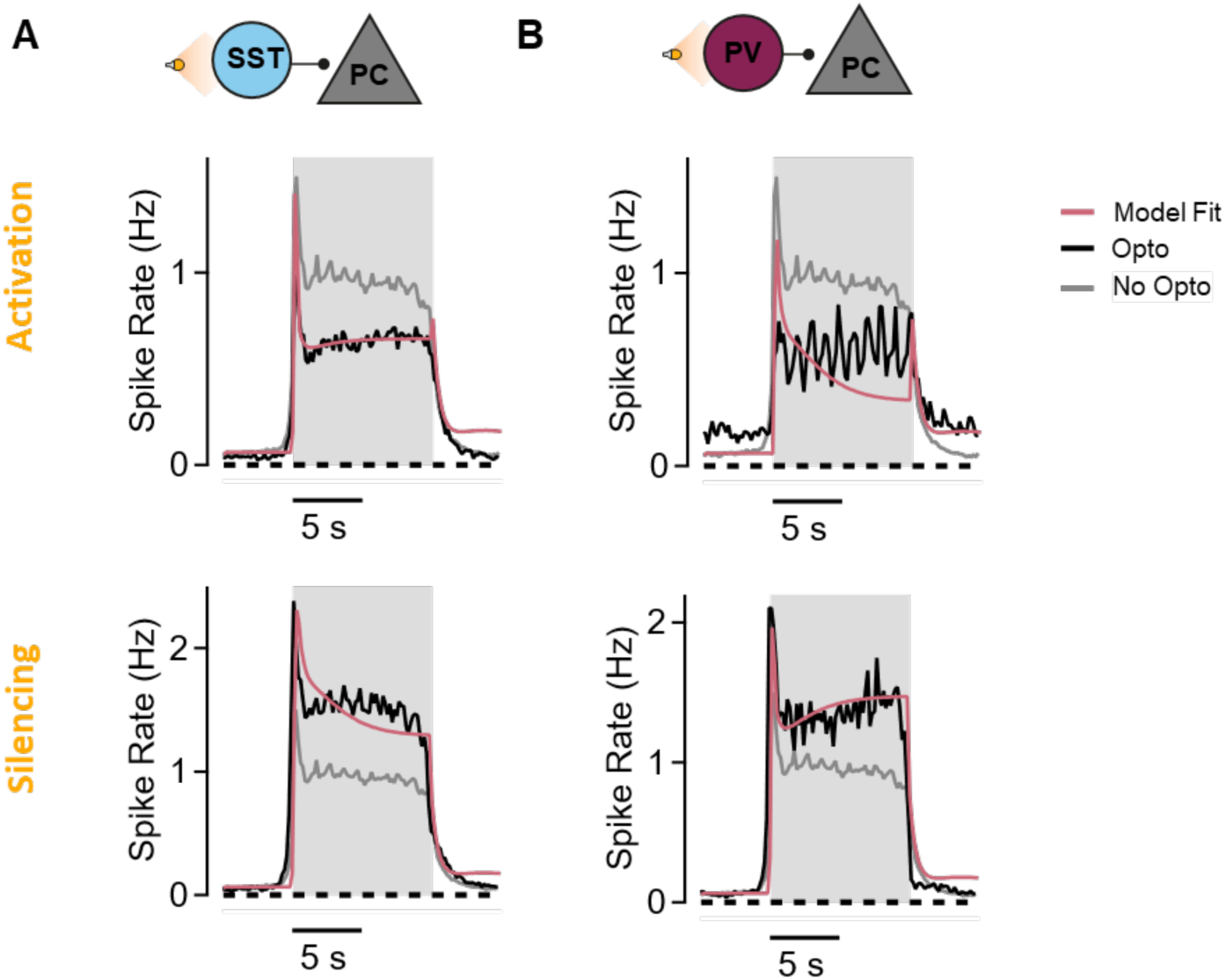
The effects of manipulating interneuron activity are predicted by the model. Average firing rates of PCs before optogenetic modulation (gray line), during optogenetics (black line) and model prediction with a change in activity of interneurons equivalent to the experimental optogenetic modulation (red line). **A**. Top: Activation of SSTs decreases PC gain and increases sensitization. Bottom: Silencing of SSTs results in an increase in gain and non-adaptive dynamics. **B.** Top: Activation of PVs results in a decrease in PC gain and non-adaptive dynamics. Bottom: Silencing PVs increases PC gain and makes them more sensitizing.

### The average inhibitory connectivity of pyramidal cells

The major motivation for counting synaptic connections in structural data is to estimate how strongly neurons influence each other during activity, a property more directly measured by electrophysiology. Considerable efforts have been made to make functional measurements of connection probabilities and connection strengths. If electrodes are placed on a PC and a nearby PV interneuron the probability of them being connected is only ∼0.37 at distances less than ∼150 μm^67–69^. The average probability of an SST interneuron is even lower at ∼0.27. These data can be used to estimate the average PV:SST input ratio if we also know how many interneurons are available within the average connection distance of a PC (Methods). Gathering this data from a number of studies, the densities of PCs, PVs and SSTs are estimated as 100,000 cells/mm^3^, 7,856 cells/mm^3^ and 6,771 cells/mm^3^, respectively ^76^, which yields an average of 34 connected PV interneurons and 17 SSTs. Taking into account that each PV synapse is 1.55 times as powerful as each SST^69^, the average PV:SST input ratio is expected to be 3.1. The model estimates synaptic weights independently of these electrophysiological and structural data but was in close agreement with an average ratio of 2.94 (Fig. 5E).

### Random inhibitory connections can account for opposite forms of adaptation

To understand the differences in local inhbition underlying opposing forms of adaptation we fit the model separately to the dynamics of each of the three subsets of PC defined functionally in Fig. 1 - sensitizers, non-adaptors and depressors (Fig.7). Two trends were observed as PCs altered from sensitization towards depression: SST inputs became weaker and PV inputs stronger (Fig. 7A right). Qualitatively, these changes can be easily understood on the basis of the average responses within each inhibitory population: PVs are sensitizing and are therefore expected to drive PCs to depression while SST inputs are depressing and therefore drive PCs to sensitization (Fig. 4A). The model now provides *quantitative* predictions of how the PV:SST balance varies to drive adptation in opposing directions: this ratio averaged 2.3 in the sensitizing tertile of PCs, 3.0 in the non-adaptors and 4.1 in the depressors (Fig. 7A).

**Figure 7.**
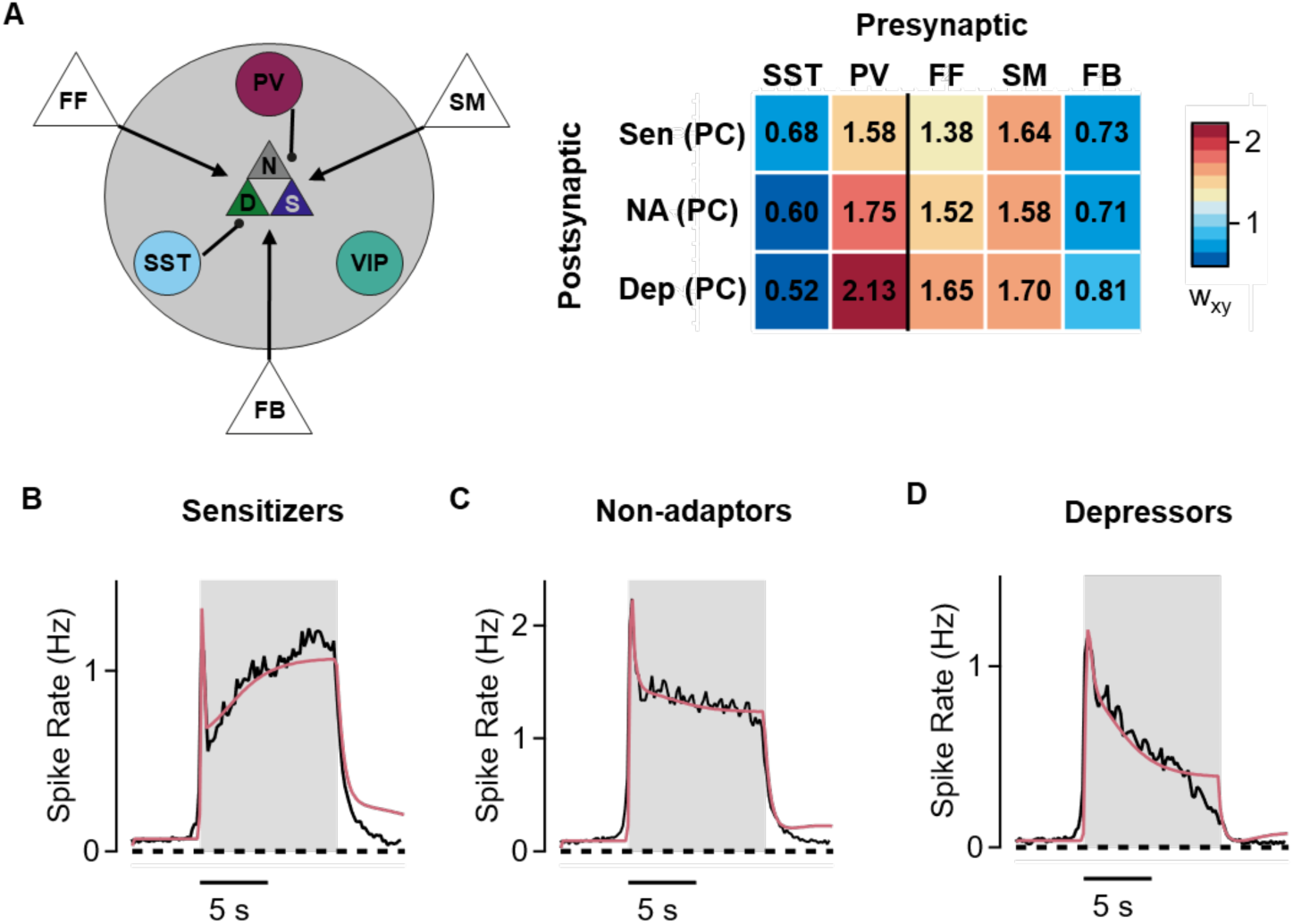
Connectivity underlying different modes of adaptation. **A**. Left: Scheme of the model used to fit separate circuits for depressors and sensitizers. We fixed all the indirect connections because it was not possible to know whether they would be different in depressors and sensitizers (connections not showed). We only let free to vary those direct presynaptic connections to PCs (black connections). Right: Heatmap of total synaptic weights (wxy). Notice the higher SST inhibition of sensitizers and the higher PV inhibition of depressors. **B.** Average firing rates (black) and model fitting (light red) of sensitizers. They receive stronger SST inhibition **C.** Same as in D of non-adaptor PCs. **D.** Same as in D of depressor PCs. They receive stronger PV inhibition.

Some variability in the strength of connections from PV and SST interneurons onto PCs is to be expected. But how much? And how does this variability arise? Are there *specific* rules based on molecular and/or functional differences that generate two general classes of microcircuit within layer 2/3, one causing PCs to depress and the other to sensitize^3,13,17^? The distribution of AIs measured experimentally was unimodal and did not, therefore, provide statistical support for this idea (Fig. 1D). We therefore began by investigating the null hypothesis - random connectivity ^8,22,24,25,77^.

Peters’ rule posits that the number of synaptic connections between two neurons depends on the geometric overlap of their dendritic trees and axonal arbors. If variations in overlap occur randomly, we can estimate the complete distribution of PV:SST input ratios from the known cell densities ^76^ and average connection probabilities^67–69,78^. This was achieved by simulating PV and SST inputs onto each of 100,000 PCs by interrogating each interneuron within 150 μm of the PC and deciding if it was connected by flipping a coin loaded to an average probability of 0.37 for a PV and 0.27 for an SST (Fig. 8A and B; see Methods). A distance of 150 μm was used because almost all connections occur within this range^67,69^. The simulated distribution was skewed towards higher PV:SST ratios because of the higher density of PV interneurons and their higher connection probabilities (Fig. 8B). The mean PV:SST ratio for the first tertile was 2.47, which compared to an estimate of 2.31 obtained by fitting the model to the dynamics of sensitizing PCs (Fig. 8C). In the third tertile of he simuated distribution the average PV:SST ratio was 3.94 compared to a value 4.08 obtained by fitting the model to the dynamics of depressing PCs. A finer comparison of the relation between AI and PV:SST input ratios was obtained by mapping values based on the structural simulation and dynamic model using 130 bins rather than just three (Fig. 8D). The relation was sigificantly steeper for sensitizing neurons (0.57 ± 0.009) compared to depressors (0.35 ± 0.002, green; p < 10^-4^, t-test).

**Figure 8.**
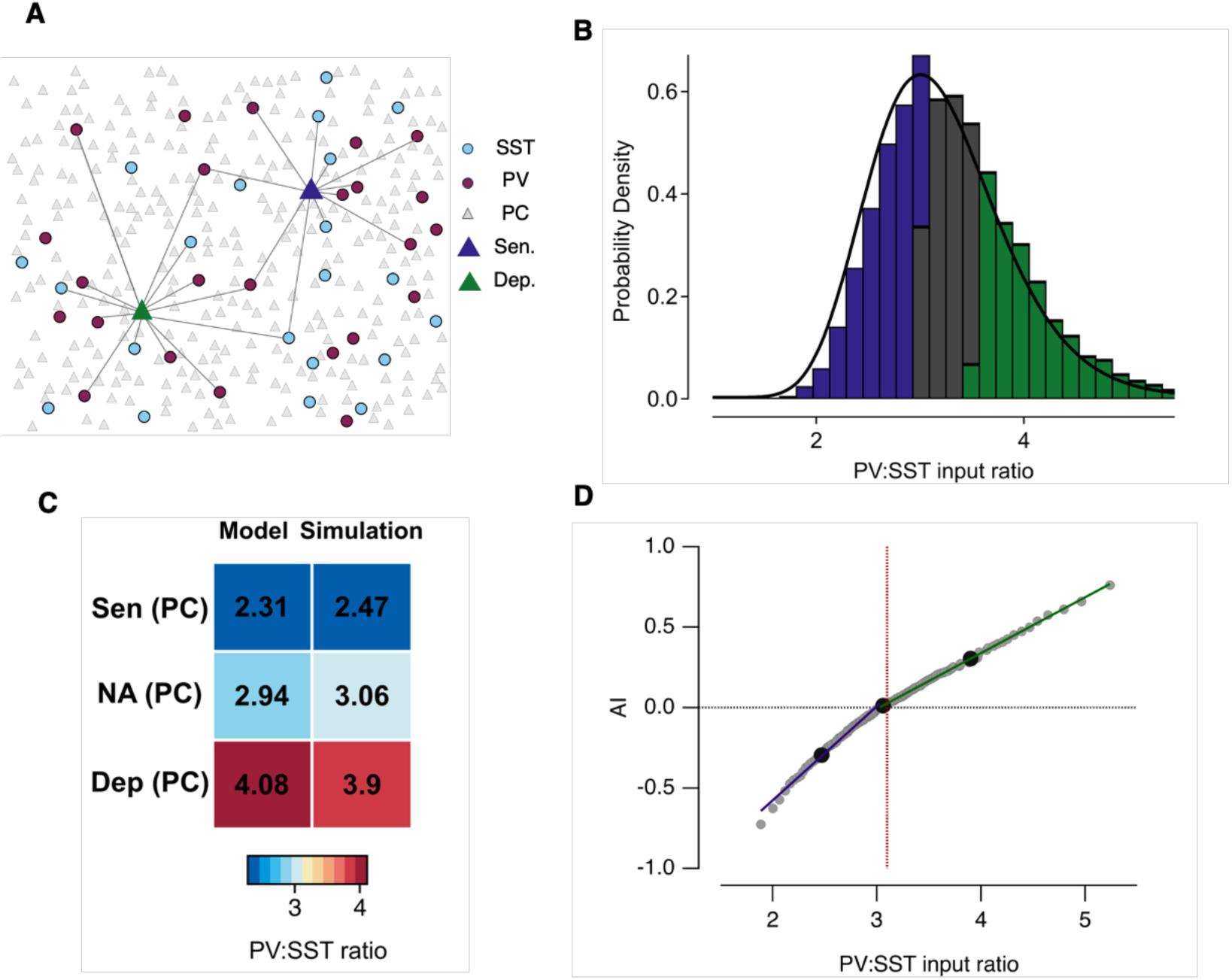
Random inhibitory connections to PCs can account for opposing forms of adaptation. **A.** Schematic of the simulation showing the relative densities of the four major types of neuron. Two PCs are highlighted with some of their connected interneurons (depressing, green; sensitizing, blue). The connection from a given interneuron to a PC was set by tossing a coin loaded accoroding to the connection probability that has been measured by electrophysiology. **B.** Distribution of the PV:SST input ratios from a simulation of 100,000 PCs was skewed towards higher ratios and described by a logNormal function with mean 3 and standard deviation 0.38. Blue bars: lower tertile corresponding to sensitizers. Green bars: upper tertile corresponding to depressors. **C**. Comparison of the PV:SST input ratios calculated from the dynamic model and from the simulation based on random connections. **D**. AI distribution mapped onto PV:SST input ratio. Large black circles show the distributions split into tertiles (data from C) and grey dots split into 130 bins. AI changes linearly with PV:SST ratio with higher slope for sensitizers (0.57 ± 0.009, blue) than for depressors (0.35 ± 0.002, green). The red line shows the average PV:SST input ratio (3.1) calculated from structural and electrophysiological data.

These results demonstare that random connectivity from PV and SSTs to PCs is sufficient to account for opposing forms of adaptation in V1 - specific connectivity rules do not need to be invoked. We conclude that the local inhibitory circuits under which PCs operate are far from functionally uniform ^5,14,16,79^. Rather, random ariations in the strength of PV and SST inputs lead to profoundly heterogenous adaptive responses in PCs, driven by slow modulatory inputs to layer 2/3.

## Discussion

The different patterns in which interneurons connect with pyramidal cells determine the varied ways in which sensory inputs are processed in the cortex ^1–7^. Here we have focused on a fundamental aspect of this issue: the relation between *variations* in local connectivity and *variations* in function. The process we analyzed, adaptation, is common to all sensory systems and we investigated it’s operation over time-scale of several seconds (Fig. 1) when long-range inputs to V1 are activated ^43,57^. The use of a circuit model that incorporates measurements of activity from the three major classes of interneuron (Figs. 2-4) provided quantitative estimates of the way in which variations in local inhbitory circuits generate depressing and sensitizating modes of adaptation in PCs (Figs. 5-7). The balance between PV and SST inputs is a key parameter because these interneuron populations adapt in opposite directions to the visual stimulus (Fig. 1). Variations in this balance could be accounted for by two general statistics of the cortical connectome operating under a random connectivity rule (Fig. 8). First, interneurons occur at much lower density than PCs ^76^. Second, an individual interneuron only contacts a small fraction of nearby PCs. These results provide a stark example of the impact of Peters’ rule on cortical processing^8,22–25^.

### “A statistical summary of the connectome”

A random connectivity rule underlying variations in adaptation is wholly compatible with the idea that specific connectivity rules also operate within layer 2/3 to determine other functions. It is clear, for instance, that SST interneurons target PCs while avoiding contact with each other ^12^, while PCs of similar tuning are more likely to be connected ^80^. A recent connectomic study of mouse visual cortex compared the number of inhibitory synapses on the soma and dendrites of PCs (roughly corresponding to PV and SST inputs) and found a correlation coefficient of ∼0.5 in layer 2/3 ^8^ leading to the conclusion that the number of PV and SST inputs are determined by a combination of spatial overlap and more specific synaptic targetting. The present work complements such connectomic data by characterizing the functional impact of the differences in local inhibition that arise randomly. Such a “statistical summary of the connectome”^24^ will be an important tool in understanding cortical function. We suggest that such a summary should include not only general circuit motifs but also the distributions that describe how these motifs vary (Fig. 8B).

There are large differences in the densities and connection probabilities of excitatory and inhbitory neurons in different areas of cortex, between layers within an area and even at different depths within a layer ^8,81^. For instance, the density of pyramidal cells in layer 2/3 of somatosensory cortex is about five times that in auditory cortex ^76,82^. There, the ratio of excitatory to inhibitory neurons is only 2:1 compared to ∼6:1 in auditory cortex. Even within layer 2/3 of visual cortex, the probability of a connection between an SST interneuron and pyramidal cell is lower in the more superficial zone ^68^. We are far from understanding the functional impact of these variations but this study indicates that the PV:SST input ratio may be a useful summary metric for understanding how variations in local inhibition alter PC function.

### A long-range signal that decorrelates pyramidal cells

The utility of summary statistics will depend on the computation being investigated and the time-scale on which it is observed. It would be interesting, for instance, to understand how the fast phase of adaptation, dominated by depression (Fig. 1), varies across the PC population and whether this also depends on the balance between PV and SST inputs. During the first 0.5 s of stimulus presentation feedforward signals carry the major external drive to layer 2/3 and depression of these synapses is a major cause of depressing adaptation in PCs ^83^. But on longer time-scales activity in V1 is also dependent on feedback signals from higher visual areas aswell as slower top-down inputs from other cortical and subcortical areas and these are crucial for computations involving behavioural goals ^52^. Here, we idenitified a powerful sensitizing input that impacts PCs both directly and though VIP and PV interneurons (Figs. 2 and 3). Most models of V1 dynamics have neglected the role of slower and long-range connections and confined themselves to considering feedforward inputs ^68,84–86^, in large part because we are only just starting to learn how slower inputs modulate visual processing^87^.

The antero-medial cortex is the most likely source of the slow modulatory signal since these inputs to V1 show sensitization in response to a moving grating stimulus ^43^. Inputs from retrosplenial cortex to layer 2/3 also sensitize during learning ^57^. as do signals from layer 4 ^88^, indicating that there may be alternative routes by which this co-ordinating signal enters layer 2/3. VIPs are a major target of long-range inputs to V1 (Fig. 3) and will increase the PV:SST input ratio through the VIP->SST->PC disinhibitory circuit. But, whatever the source(s) of the slow modulatory signal, it’s basic effect is to decorrelate activity across the population of PCs because of variations in their PV:SST input ratios (Fig. 1A-E).

Sensitization becomes the dominant form of contrast adaptation when a mouse is conditioned to a reward but the dominant form of adaptation is depressing when the stimulus is not behaviourally relevant ^89^. This leads to the suggestion that the balance between sensitizing and depressing adaptation is modulated by top-down inputs to enhance the signalling of stimuli that have become behaviourally important. It will be interesting to understand whether such simple forms of learning involves changes in the PV:SST input ratio or other aspects of the circuits in layer 2/3. The model that we have applied to understand the circuits underlying adaptation will provide a tool for investigating changes underlying processes such as habituation or reward-association.

## Methods

### Animal preparation and multiphoton imaging *in vivo*

All experimental procedures in mice were conducted according to the UK Animals Scientific Procedures Act (1986). These were performed at University of Sussex under personal and project licenses granted by the Home Office following review by the Animal Welfare and Ethics Review Board.

Experiments were performed with adult mice (P60-P90) and were described in detail previously ^42^. Results are reported from eighteen SST-Cre mice (SST tm2.1(cre)Zjh/J, Jackson #013044), ten PV-Cre (Pvalb tm1(cre)Arbr/J, Jackson #008069), and four VIP-Cre (VIP tm1(cre)Zjh/J Jackson #010908) from either sex. Mice were prepared for head-fixed multiphoton imaging through a surgical procedure in a stereotaxic frame under isoflurane anaesthesia. A titanium head plate was attached to the skull and a 3 mm craniotomy drilled to expose the brain. We then expressed AAV1.CaMKII.GCaMP6f.WPRE.SV40 to image calcium activity in pyramidal cells and AAV9.CAG.Flex.GCaMP6f.WPRE.SV40 to image interneurons in Cre lines. To excite interneurons optogenetically we used rAAV9/Syn.Flex.ChrimsonR.tdTomato and to inhibit we used AAV5.CBA.Flex.ArchT-tdTomato.WPRE.SV40.

During imaging mice were free to run on a cylindrical treadmill placed under a two-photon microscope, with stimuli being presented io two monitors covering most of the visual field. Visual stimuli were sinusoidal gratings drifting upwards (100% contrast, 1 Hz, 10 s duration) generated using PsychoPy ^90^. The amplitude of visual responses in V1 are strongly modulated by locomotor activity and here we have focused on the running state, only using mice that were naïve to the stimulus to avoid any effects of habituation. The standard experimental protocol consisted of ten presentations of the stimulus with a 20 s interstimulus interval. When testing optogenetic manipulations these 10 trials were interleaved with 10 additional trials paired with illumination from an amber LED. The intensity of this LED was tuned to modify the activity of PCs to be double when silencing interneurons with ArchT or half when activating interneurons with ChrimsonR. Running speed was measured with a rotary encoder and directly fed into the two-photon imaging software (Scanimage5; Vidrio Technologies). The two-photon microscope (Scientifica SP1) employed a 16x water immersion objective (0.8 NA; Nikon) and imaging was carried out at a depth of 150-300 μm below the surface of the brain.

Raw movies were registered and segmented into regions of interest (ROIs) using the Suite2P package running in MATLAB 2021a^91^ and further analysed using custom-written code in Igor Pro 8 (Wavemetrics). Here we only analyse results from the first session in which the animal was exposed to the stimulus to avoid changes in activity and adaptive properties reflecting habituation to the stimulus. Stimulus trials were only included for analysis if the mouse was running faster than 3 cm/s for more than 80% of the time. To calculate the response dynamics of PC, PV, SST and VIP populations we only included cells that were positively correlated with the stimulus.

### Estimation of firing rates

To estimate firing rates from the recorded ΔF/F data we used the MLspike algorithm using initial parameters expected for GCaMP6f ^46^. Briefly, this finds the optimal spike train to fit the calcium fluorescence trace (ΔF/F) taking into acount estimates of baseline fluorescence level and neuronal parameters including the amplitude of the unitary Ca^2+^ transient generated by a spike (A) and the decay time-constant of that transient (τ). In general, we followed recommendations and demos in the article and on the MLspike GitHub page. Some initial parameters were obtained from (Deneux et al., 2016). We set the range for A to between 1 and 50% and τ was set at values from 0.05 to 3 seconds. The parameter Amax, which corresponds to the maximum amplitude of detected events, was 9, and the saturation parameter was set to 0.002. The Hill coefficient for Ca^2+^ binding to GCaMP6f ws free in the range from 1 to 3.75 with step sizes of 0.25.

### Model of the circuit in layer 2/3 of V1

We built rate-based population model of the network circuit in the mouse visual cortex layer II/III. The model was aimed to describe and predict the behaviour and interconnection of four main neuron populations (PC, PV, SST, and VIP) during processes of adaptation, as well as the effect of optogenetic manipulation. The network was represented by the system of four mean-field first order ordinary differential equations established by Wilson and Cowan ^74^:

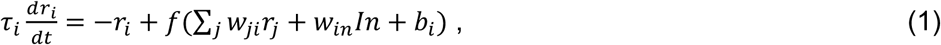

where *i*, *j* are iterating through populations, *r* is a firing rate. Time constants *τ_PC_* = 15 *ms*, *τ_pV_* = 7.5 *ms*, *τ_SST_* = 19 *ms*, *τ_VIP_* = 19 *ms* were set according to other work ^92^. *f*(*x*) is a neuronal input-output function and in this work we used a rectified power low ^93^):

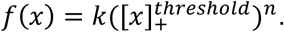

Here, *k* is a scaling coefficient which was set to 1. The dynamics of the system reaches supralinear form over the same range of firing rates as previous studies ^85,88^. 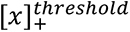 corresponds to rectified linear function with a lower threshold set to 0 to prevent negative values of firing rate, and upper threshold set to 25 – an arbitrary value set to prevent run-away behaviour of the algorithm solving the network equations system. The exponent value was set to *n* = 2. The linear activation function (*k* = 1, *n* = 1) gave similar results during the fitting process^94^.

Connectivity among populations were represented by synaptic weights *w_ji_* (Equation 1) where *j* is a presynaptic population and *i* postsynaptic. The total synaptic weight between populations *j* and *i* is a compound parameter of synapse properties:

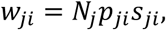

where *i*, *j* are iterating through populations, *N_j_* is the number of responsive presynaptic cells, *p_ji_* is an average connection probability between populations and *s_ji_* is the average synaptic strength. The model operates in terms of *total* synaptic weights, dependent on the number of responsive cells in the presynaptic population. To obtain the average synaptic weight, the total synaptic weight was divided by the number of responsive presynaptic cells.

The aim of fitting the model to the average activity traces of neuronal populations was to obtain synaptic weights proportional to the number and strength of synaptic connections between populations. Where experiments indicate that connections are very weak or non-existent, as between SST interneurons^12,68,70^, weights were set to zero.

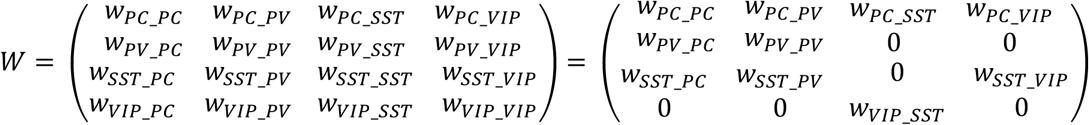

#### External inputs driving activity

Three types of external input drove activity in the circuit: feedforward (FF), slow modulation (SM) and feedback input (FB), the forms of which are discussed in Results. These were represented by a general variable *In* that could differ for each population. The feedforward input was defined by the following system of equations:

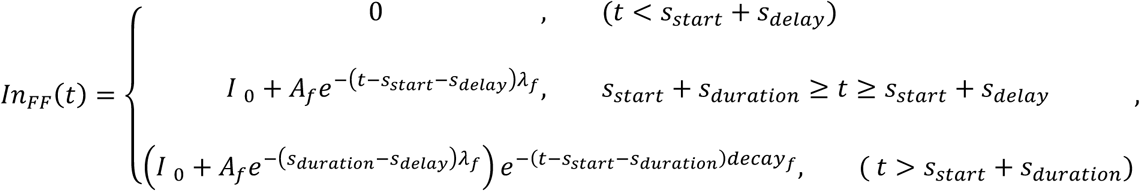

where *s_start_*, *s_delay_*, *s_duration_* represent stimulus start, delay in the initation of the response in a cell and the stimulus duration, respectively. *I_0_* is a base activity that was set to *1 Hz* and represents a step input component after short fast-depressing component, defined by exponential decay with an amplitude *A_f_* and decay rate *λ_f_*. At end of the stimulus representation, experimental traces decayed relatively slowly compared with the time-constant of the model neurons and this was accounted for by incuding an exponentially decaying component to the input with the argument *decay_f_*.

The temporal form of the feedforward input measured experimentally reflects application of spatio-temporal Gabor filter to a sinusoidal wave of drifting gratings^95^. This oscillation was clear in the activity of individual PCs but much less clear in population averages because neurons responded with different phases, depending on the location of their receptive fields relative to the stimulus location. This 1 Hz oscillations were most clearly observed in the average response of the SST population (Fig. 1F) which have bigger receptive fields and therefore display less phase shifting ^64,96^. We therefore did not attempt to model the oscillatory component of individual neuronal responses and concentrated on capturing the average activity across populations. FF was therefore defined as a step of duration 𝑠_𝑑𝑢*r*𝑎𝑡*i*𝑜*n*_with initial amplitude (*I_0_* + *A_f_*) which then decayed to a steady state of *I_0_* with time-constant 210 ms with these parameters being obtained from fitting initial peak in PCs data (Fig. 1C). The steady state was half the initial maximum with *A_f_* = *I_0_*. The feedforward input from layer 4 to layer 2/3 displayes two time-constants of depression, with the slower of the order of 10 s^97^ but including this slower component of depression failed to account for the slower dynamics of our data and was therefore omitted from the model.

The slow modulatory input was represented by the following system of equations:

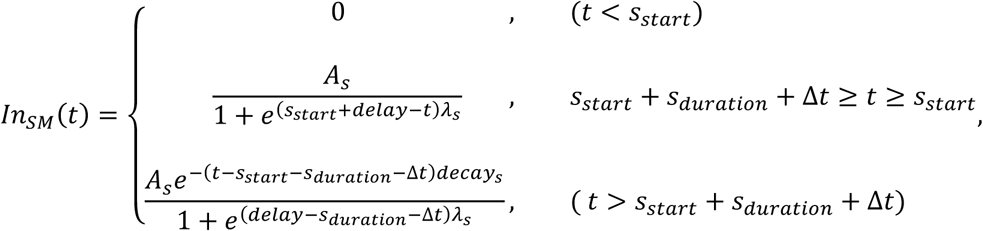

where 𝑠_𝑠𝑡𝑎*r*𝑡_, 𝑠_𝑑𝑢*r*𝑎𝑡*i*𝑜*n*_ represent time of the stimulus start and the stimulus duration, respectively. The slow component is represented by a sigmoid function with an amplitude 𝐴_𝑠_, growth rate 𝜆_𝑠_, and centre of sigmoid shifted to a 𝑑𝑒𝑙𝑎𝑦 value from the stimulus starting time.

Δ𝑡=0.165 s. This delay was introduced based on the observation of a small (one sample point) delay between the end of the stimulus and the beginning of response decay in PV and VIP cells. Again the decay phase was accounted for by the argument 𝑑𝑒𝑐𝑎𝑦_𝑠_.

Finally, feedback input was represented by a step function:

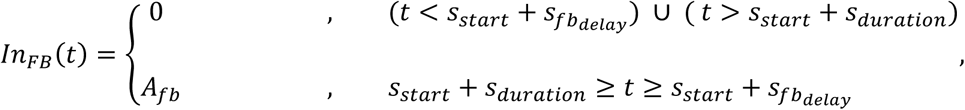

where *s_start_*, *s_fb_delay_*, *s_duration_* represent time of the stimulus start, delay in the initation of the response in a cell and the stimulus duration, respectively. The amplitude of the stimulus is defined by *A_fb_*. The amplitude of the stimulus is defined by *A_fb_*. In the three external inputs, the amplitudes (𝐴_𝑓𝑏_, 𝐴_𝑠_, *I_0_*) were fixed around 1 Hz based on the average response of PCs (Fig. 1C).

### Fitting the model

To obtain values of synaptic weights we fit the model simultaneously to the average activity traces of the four neuronal populations using the *LMFIT* library in Python^98^ by minimization of reduced chi-square parameter:

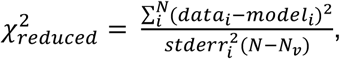

where (𝑑𝑎𝑡𝑎_*i*_ − 𝑚𝑜𝑑𝑒𝑙_*i*_)/ 𝑠𝑡𝑑𝑒*rr*_*i*_ represents the residual function weighted by uncertainties in the experimental data, *N* is the number of data points and *N*_𝑣_ is the number of variable parameters. We set the threshold chi-square value to be 8,.

The model incorporates about 20 parameters free to vary and we compared several fitting methods: *Levenberg-Marquardt, Nelder-Mead, Powell, L-BFGS-B*. We noticed that the *Nelder-Mead and Powell* methods worked well on the full range of values defined by minimum and maximum limits providing initial parameters for which *Levenberg-Marquardt* and *L-BFGS-B* provided more precise fits.

Different sets of parameters sometimes provided similar values of reduced chi-square in multidimensional parameter space. An important further constraint was provided by requiring the model to additionally fit the results of optogenetic manipulations of the PV and SST populations (Fig. 6), leading to a reduced number of sets of possible synaptic weights. The intensity of the LED was set to increase or decrease PV or SST activity by a factor of 2 (as measured empirically in other parallel experiments measuring GCaMP6f responses in these interneurons^42^). The average PV or SST activity trace was then scaled by the same factor of 2 for fitting the model to observed PC actvity. This approch allowed us to find a stable set of parameters describing both the basic state of the circuit and the effects of perturbing the circuit.

### Comparing the model to other studies

We compared the value of synaptic weights from our model to connectivity measures made using electrophysiology ^68,69^ and another model^75^ (Figure 5F). A simple correction was made to allow this comparison. Our model was fit to the average activity of responsive cells only, while these other studies obtained average synaptic weights by multiplying the connection probabilities and synaptic strengths of the whole presynaptic population. The synaptic weights from the model were therefore divided by the number of responsive cells per field of view of each presynaptic population to obtain what we have termed average synaptic weights (PC = 40 Cells/FOV, SST = 11 Cells/FOV, PV= 30 Cells/FOV).

### Fitting the model to subpopulations of pyramidal cells

Sensitizers, non-adaptors and depressors were fit separately to assess the differences in their connectivity. The population of PCs was separated in three tertiles based on their AI and the response of these three groups was averaged (Fig. 7). The model was then fit with three simplifying assumptions or steps. First, that each subpopulation of PC receives the same shape of inputs as the average across the population. Second, only the synaptic weight of those connections directly targeting PCs were left free to vary while the other connections within the circuit were fixed. Third, we used the initial conditions of the average model, and the final synaptic weights were corrected based on the amplitude of the average response of each tertile. Corresponding multiplication factors are sensitizers = 1.015, non-adaptors = 1.182, depressors = 0.748.

### Simulation of PV:SST input ratio based on random connectivity

Because the density of PC cells is higher than PV and SST, and because probability of connection from PV to PC and from SST to PC is lower than 1 there should be a broad distribution of the number of connected interneurons to PC and therefore distribution of proportional inhibition of PC cells by PV and SST. To test an assumption that randomness in connectivity between PV to PC and SST to PC cells can describe different polarities of slow adaptation observed in this work, we simulated 100000 PC cells and analysed a distribution of ratio of PV=>PC inhibition to SST=>PC inhibition.

To be unbiased to the model results, we gathered all required information from the literature. Probabilities of connections from interneurons to pyramidal cells were set to *p*_𝑃𝑉_ = 0.375 and *p*_𝑆𝑆𝑇_ = 0.275 based on experimental data ^67–69^. The average number PV and SST interneurons within 150 μm of each PC was 111 and 95, respectively, simply calculated from the measured densities of PCs (100,000 cells/mm^3),^ PVs (7856 cells/mm^3^) and SSTs (6771 cells/mm^3^) in layer 2/3^76^. The strength of an individual synaptic connection was based on measurements of the unitary post-synaptic potential of 𝑠_𝑃𝑉_ = 0.48 mV and 𝑠_𝑆𝑆𝑇_ = 0.31 mV ^69^.

Simulation for each pyramidal cell was performed in Python by drawing a samples from a binomial distribution, represented as *n*𝑢𝑚*p*𝑦. *r*𝑎*n*𝑑𝑜𝑚. 𝑏*in*𝑜𝑚*i*𝑎𝑙(*n*, *p*, 𝑠*i*𝑧𝑒) in numpy library, where *n* = 1 corresponds to number of trials, *p* - probability of connection, 𝑠*i*𝑧𝑒 - number of inhibitory cells. This is equivalent to tossing a coin with weights of p (connection) and 1-p (no connection) on either side. The coin is tossed 1 time for each interneuron around the picked pyramidal cell. The total ratio of PV:SST inhibition of PC cells is:

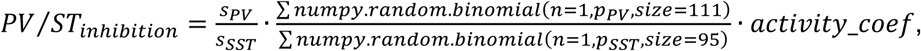

where 𝑠_𝑃𝑉_, 𝑠_𝑆𝑆𝑇_ are average strength of synapses and 𝑎𝑐𝑡*i*𝑣*i*𝑡𝑦_𝑐𝑜𝑒𝑓 = 1.25 is the ratio of proportion of active PV and SST cells (0.81 and 0.65, respectively). The simulated distribution of PV:SST input ratios was skewed reflecting the higher density and connection probabilities of PV interneurons compared to SSTs (Fig. 8B).

## Acknowledgements

This work was supported by BBSRC grant BB/X009386/1, a Ph.D scholarship to SD from the Leverhulme Trust (DS-2017-011) and a Researchers at Risk Fellowship to YK from the British Academy (RaR/100503) .

---

**Supplementary Figure 1.**
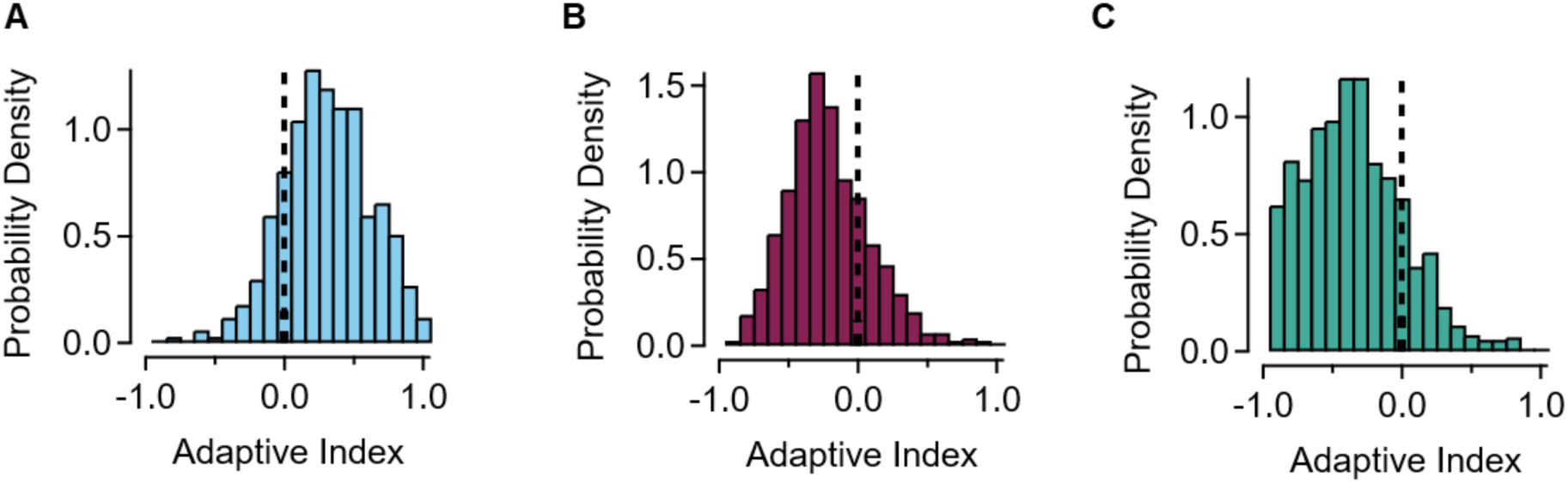
Distribution of adaptive indices in inhibitory cell populations. **A**. SST’s, showing that the large majority most of them are depressing. **B.** PV’s showing that the large majority are sensitizing. **C.** VIPs, showing that the large majority are sensitizing.

**Supplementary Figure 2.**
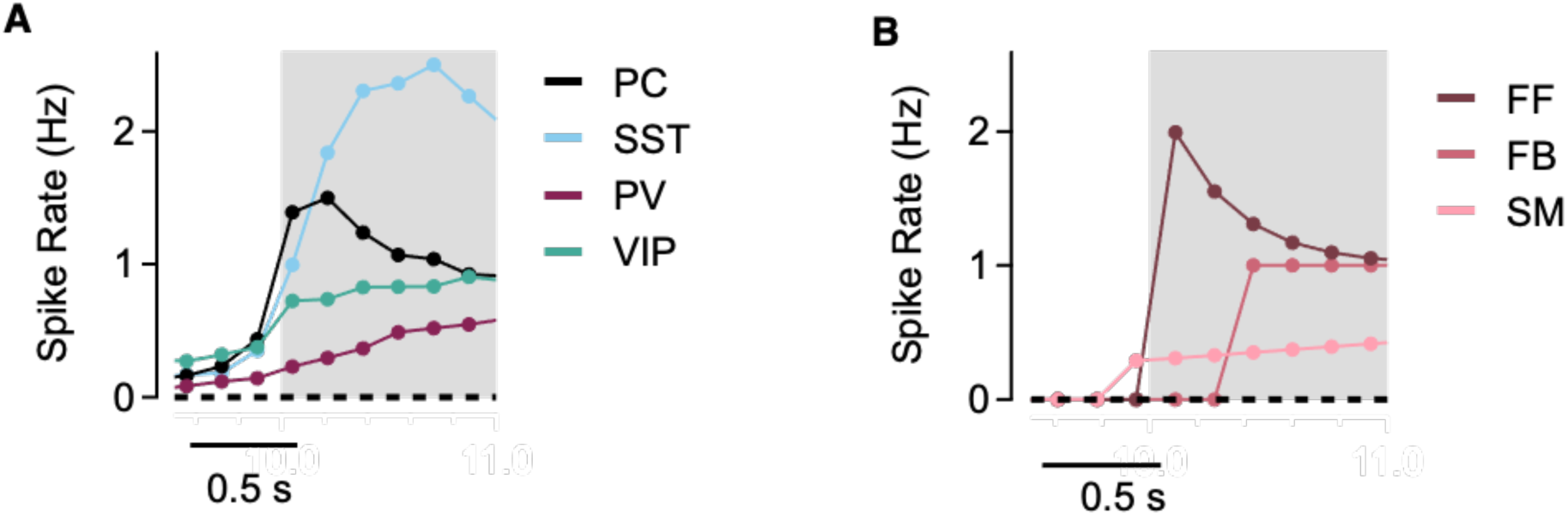
Delay of responses and external inputs relative to stimulus start. **A**. The first 1 s of the average response of each neuronal population to the high-contrast stimulus (grey bar). PCs peak first (∼150 ms) but the peak in SSTs is delayed roughly by ∼350 ms. The model could not reproduce this feature without assuming that the feedback signal (FB) was delayed relative to the feedforward (FF). **B.** Delays to peak of external inputs required to describe the differing delays in the peak of responses in A. FF (Feedforward) input peaks 212 ms after stimulus start; FB (Feedback) input 379 ms; SM (Slow modulation) input 10 s (see Fig. 2).

